# Optimization of citric acid production from sugarcane molasses using *Aspergillus niger* by submerged Fermentation

**DOI:** 10.1101/2024.10.15.618342

**Authors:** Al Amin, Hd. Razu Ahmmed, Mohammad Ismail, Tajreen Naziba Islam, Mohammed Mohasin

## Abstract

The potentiality of citric acid on economy is high because of its multi-purpose uses, particularly in the food and pharmaceutical industries. Bangladesh spent more than million US dollars to import citric acid mostly from India and China. Its consumption is increasing 3.5–4%, annually, indicating the need for better manufacturing alternatives. Globally, citric acid is primarily produced through microbial fermentation with *Aspergillus niger*. To support the massive scale of production of citrate, the manufacturing process must be eco-friendly which should be inexpensive and available raw materials for maintaining high yielding in a cost-effective manner. In Bangladesh prospective, the current study has undertaken to optimize citric acid production using one of the most abundant raw materials sugarcane molasses. Moreover, the aim of this study was to determine the optimum conditions to produce citric acid from sugarcane molasses using *Aspergillus niger (F-81)* by submerged fermentation. The amount of citric acid production was determined by Marier-Boulet colorimetric method. The optimization data suggested that 10% substrate (from processed cane molasses), 4% inoculum size of *A. niger*, and initial pH 6.0 allowed to produce around 25.8 g/L citric acid. Further study is warranted to assess the feasibility of citrate production in an industrial level as well as improvement of microbial strains is needed to further enhance citric acid production.

## Introduction

Citric acid is one of the most important tricarboxylic organic and water-soluble acids that has both food-additive and pharmaceutical values. In early twentieth century, citric acid was mostly produced by citrus rich food such as lemon and lime juices (King & Cheetham, 1987). Later on, *Aspergillus niger* was introduced to produce citric acid commercially in 1923 (Rohr, 1983). In Bangladesh, sugarcane is the second largest cash crops (Rahman et al., 2016) and approximately four million metric tons of sugarcane was produced in 2016-2017 (BBS, 2018). Moreover, sugarcane molasses is a rich source of carbohydrates for citric acid production (GUPTA & SHARMA, 1994). In addition, sugarcane contributes to produce around 0.20 million tones of sugar and 0.60 million tones molasses in Bangladesh (BSRI, 2019). Although Bangladesh has sufficient raw materials (e.g. sugarcane molasses), infra-structural setup to produce citric acid in the industrial level to meet their own demand, this year it spent more than one million US dollar to import 448 shipments of citric acid from India and China (Volza, 2024). To address this market demand of citric acid in Bangladesh, the current study has undertaken to optimize maximum production of citric acid by *A. niger* using one of the most abundant raw materials sugarcane molasses. Current study demonstrated optimum inoculum of *A. niger*, nutritional composition of cane molasses, abiotic factors like metals, pH and temperature influenced submerged fermentation-based citric acid production.

## Methods and materials

### *A. niger* growth and cane molasses preparation for submerged fermentation

The fungal strain *Aspergillus niger* F-81(91) was gifted by the “Molecular Biology Lab” of Department of Biochemistry and Molecular Biology, University of Dhaka. The spores of *Aspergillus niger* was grown on Potato Dextrose Agar (PDA) plate and incubated at 28°C for 7 days. Besides, *A. niger* also grow in Potato Dextrose Broth (PDB) a liquid media and incubated 28°C with 160 rpm and growth rate was recorded at 24 hours interval at 240 nm.

### Determination of moisture, ash, and sugar content in cane molasses

Raw cane molasses wet weight was measured and crucified, heated at 106°C for 30 minutes. Subsequently, the dry weight was measured, and the moisture content was measured by following equation (% moisture content = (Crucified weight – Dry weight)/Initial weight X 100). Besides, molasses was placed in a furnace and heated at 800°C for 2 hours. Next, dry weight was measured, and the ash content was measured as % Ash = [(weight of ash) – (crucible weight)] x 100/ [(Total crucible and sample weight) – (Crucible weight). Moreover, the sugar content was determined by comparing with standard maltose solution titration curve. Briefly, standard titration curve was determined using 3,5-dinitrosalicylic acid (DNS) which change color from yellow to red/orange in the presence of maltose. To determine the sugar content in molasses, 20 µ
L of 1M HCl was added in 1 mL of molasses solution and incubated at 90ºC for 5 minutes. Subsequently, 50 µL KOH (5N) was added to neutralize the acidic condition. The intensity of dark orange-red color was recorded at 540 nm using UV-VIS Spectrophotometer as described (Krukowski et al., 2017). Finally, the amount of sugar was measured comparing with the maltose standard curve. The pH of sample solution was recorded using a pH meter (HANNA, Germany).

### Preparation of cane molasses and fungal inoculum

Raw cane molasses was either directly prepared in distilled water without pretreatment or pretreated with potassium ferrocyanide (K4Fe(CN)6) at 90°C for 15 min. Then, ammonium salt (NH4NO3) and potassium dihydrogen phosphate (KH2PO4) were added in the molasses solution, which act as nitrogen and phosphate source, respectively. The pH was measured and adjusted to 5.0-5.6. Finally, molasses solution was sterile at 121°C under 15 psi for 20 minutes. Besides, suspension of spores was prepared from a 8 day old *A. niger* culture plate and the spores were counted by Haemocytometer (Neubauer chamber) under a light microscope and the inoculum was adjusted to 1-5 million spores/mL. Finally, the spore suspension was quantitatively inoculated into molasses solution.

### Different Fermentation conditions for citric acid production

*A. niger* spores were suspended in molasses media was placed into a shaking incubator for fermentation processes. Variations in fermentation conditions were maintained to observe the effect of different physical parameters on citric acid production. 10 ml and 15 mL of untreated/pretreated cane molasses was taken into three 500ml Erlenmeyer conical flasks and diluted by adding 190 mL/185mL distilled water to make a substrate concentration of 5% and 7.5%, respectively. Similarly, to determine optimum concentration of molasses, 10.5% and 12.5% substrate were also prepared. Initial pH of the media was adjusted to 5.5. The media was then autoclaved at 121°C under 15 psi for 20 minutes. Subsequently, 2% spore suspension was inoculated in 5-12.5% cane molasses containing solution, and incubated at 28°C, 160 rpm for 13-14 days. Similarly, to determine optimum inoculum, 1%, 2%, 4% and 8% spore suspensions were also inoculated into a 10% substrate solution and incubated 12 days at 28°C, 160 rpm to observe the citric acid production. In addition, the optimum pH was also determined using 2.0, 4.0. 6.0 and 8.0 pH adjusted 10% molasses containing substrate media and 4% spore suspension as a source inoculum and incubated for 12 days at 28°C, 160 rpm.

### Precipitation and estimation of Citrate

Fermentation broth was filtered through Whattman filter paper to remove fungal micelles and suspended materials. Then, equal amount of 10% Ca(OH)2 was added to the medium and heated for 1.30-2.0 hours at 60°C -70°C. Lime was then precipitated into tricalcium citrate tetrahydrate and the precipitate was filtered & washed with water. Thereafter, an equal amount of 20% H2SO4 was added and heated for 1.30-2.0 hours at 60°C. Next, it was filtered, and mother liquor was collected. The mother liquor was further filtered through 0.2μm PTFE-Syringe filter (Sartorius company) and the estimation of citrate was done using Marier-Boulet method (Pyridine-Acetic Anhydride method) (Alhadithy, 2020). The diluted (100 times) and filtrated mother liquor was treated with pyridine (MERCK), and then acetic anhydride (MERCK) was added and incubated 32°C for 30 minutes. Finally, the intensity of yellow color was measured at 405nm.The amount of citrate was measured through comparing with a standard calibration curve of citrate. Briefly, a stock citrate solution of 5 mg/mL was prepared by using tri-sodium-citrate dihydrate (MERCK). Then, 50-300μL of citrate solution were taken in separate test tubes and 4.95-4.70 mL distilled water was added to each test tubes, respectively. After that, 1 mL of each diluent was taken into separate test tubes including a tube containing 1 mL distilled water as blank. Next, 1.30 mL of pyridine was added to each tube and mixed thoroughly. next, 5.70 mL of acetic anhydride was added to each tube and placed the sample in water bath at constant temperature of 32°C for 30 minutes. The optical density was recorded at 405nm. Finally, a standard calibration curve was drawn taking the citric acid concentration at X-axis and optical density at Y-axis. The amount of Citrate was measured in mg/mL using the following equation. Concentration of Citrate((μg/mL) x Dilution Factor/1000.

### Statistical Analysis

All statistical analysis were done using a software called GraphPad Prism version 8.0.2. Data were expressed as mean ± SD (Standard deviation). All experiments were performed at least three replicates.

## Results

### Preparation of *A. niger* inoculum and cane molasses suspension for submerged culture

It is well-established that the branches mycelium of acidogenic *A. niger* contains short hyphae with swollen tips, which are suitable for citric acid production irrespective of their pellets or filamentous forms (Babitha et al., 2007; Snell, 1951). Moreover, the main substrates for citric acid production are solutions of sucrose, molasses or glucose (Domínguez, 2010). Previous study suggested that raw-material which contains 14-22% sugar is sufficient for fermentation-based citric acid production (M. Hossain, 1984). Visible and filamentous forms of *Aspergillus niger* F-81 spores were grown in 10 days in potato dextrose agar. Similarly, the suspension culture of *A. niger* was grown to the highest level by 10 days in a potato dextrose broth media. As source of sugar, cane-molasses were processed and crushed to a solution. The physico-chemical properties of cane-molasses suggested that it has 53.5% sugar residue, 24.7% moisture, 16.3% ash and pH was acidic (5.6).

### Determination of optimum concentration of substrate, microbial inoculum, and pH for fermentation-based citrate production

The effect of substrate concentration on citric acid production was determined at 5%, 7.5%, 10% and 12.5% substrate concentration (V/V) with certain inoculum size (2%) of *A. niger*, initial pH (5.6), and 10 days cultivation at ambient temperature (circa 28°C) with 160 rotation per minute (rpm). The maximum yield of citric acid has been recorded 12.09 mg/mL with 10% substrate concentration (V/V). Moreover, citric Acid production has also recorded 5.18 mg/mL, 6.36 mg/mL, and 7.48 mg/mL with a 5%, 7.5% and 12.5% substrate concentration, respectively (Figure 2B). Whereas the effect of fugal inoculum size was determined using 1%, 2%, 4% and 8% inoculum sizes with a certain amount of substrate concentration 5%, initial pH 5.6- and 12-days cultivation setup at an ambient temperature and 160 rpm. The maximum yield of citric acid was 17.67 mg/mL in 4% inoculum size of *A. niger*. Besides, 1%, 2% and 8% inoculum of *A. niger* spores allow to yield 3.76 mg/mL, 10.80 mg/mL, and 13.98 mg/mL of citrate, respectively (Figure 2C). In addition, pH optimization data shows that the maximum yield of citric acid was 9.70 mg/mL at initial pH 6, whereas the citrate yielding were 3.24 mg/mL, 6.62 mg/mL, and 5.85 mg/mL at pH 2, 4, and 8, respectively (Figure 2D).

**Figure 1:**
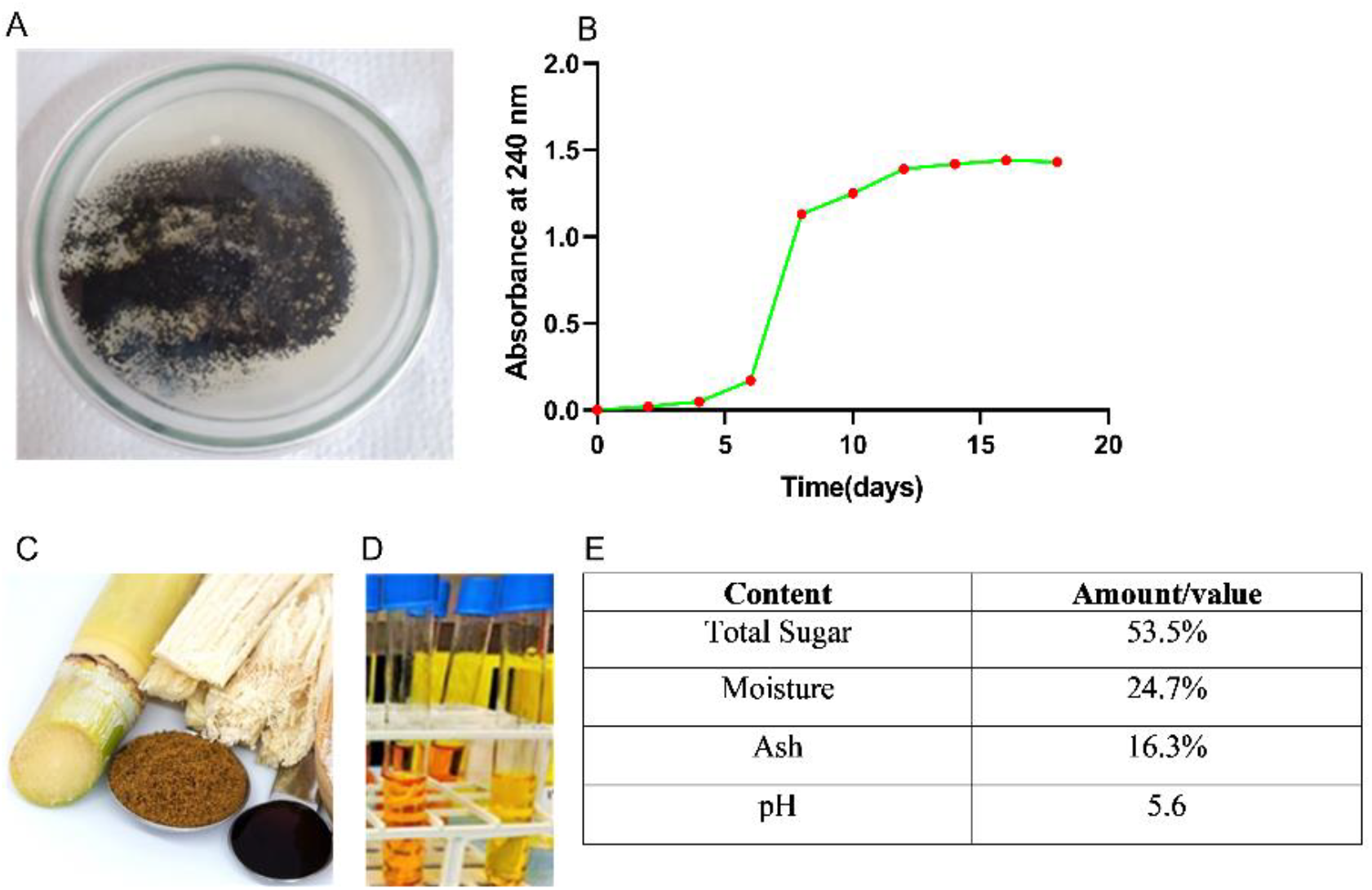
Optimization of *Aspergillus niger* growth in potato dextrose agar/broth. Figure A shows formation of fungal (*A. niger*) spores after 10 days culture on potato dextrose agar at 28°C. (B) shows the growth curve of *A. niger* in potato dextrose broth. (C) shows the raw and processed cane molasses (D) shows the maltose color reagent which was used to measure sugar content in cane molasses. Table (E) shows the physico-chemical properties of cane-molasses which were assessed by appropriate methods.

**Figure 2:**
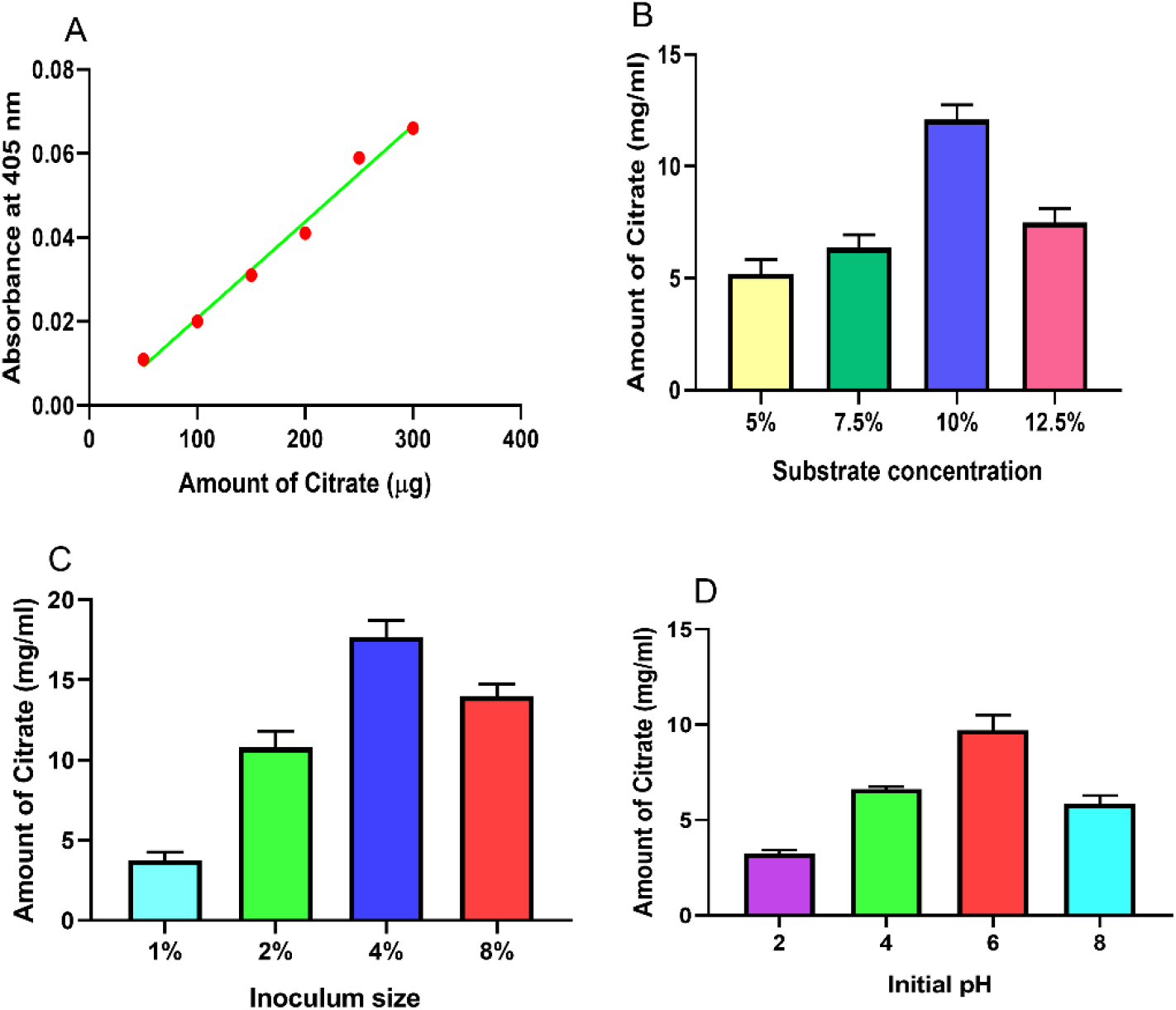
Optimization of molasses concentration, fungal inoculum, and physico-chemical features of submerged fermentation. (A) shows the titration data of citrate, which were generated by Marier-Boulet method using a standard citrate solution. (B) shows the amount of citrate production using different concentrations of molasses as a source of substrate. (C) shows the amount of citrate production using different sizes of inoculum of A. niger. And (D) shows the amount of citrate production at different pH of broth. Data shows as mean ± standard deviation. N=3.

### Citric acid production from different sugar concentration by *A. niger*

*Aspergillus niger* was first introduced to produce fermentation-based citric acid in 1919 (Papagianni, 2007). To produce citric acid (CA) by submerged fermentation process, *A. niger* inoculum spores (2% or 1.47×10^6^ spore/mL) were prepared from potato dextrose agar and sugarcane molasses were utilized as a source of cheap raw material. To determine citric acid production capacity from sugarcane molasses, a titration curve generated by Marier-Boulet method (colorimetric) using standard citrate solution (Figure 3A). The amount of citrate production in different batches of fermenter were 5.2 mg/mL to 6.2 mg/mL in 5% substrate and 6.4 mg/mL to 7.8 mg/mL in 7.5% substrate containing with 8 days fermentation (Figure 3B and 3C). As expected, there was no significant difference of citrate among the batches of fermenter. Moreover, the continuous-batch fermentation with 5% substrate of molasses, 2% inoculum of *A. niger* and initial pH 5.6 has shown 16.0 mg/mL (circa) citrate has yielded by 16 days of cultivation (Figure 3D).

**Figure 3:**
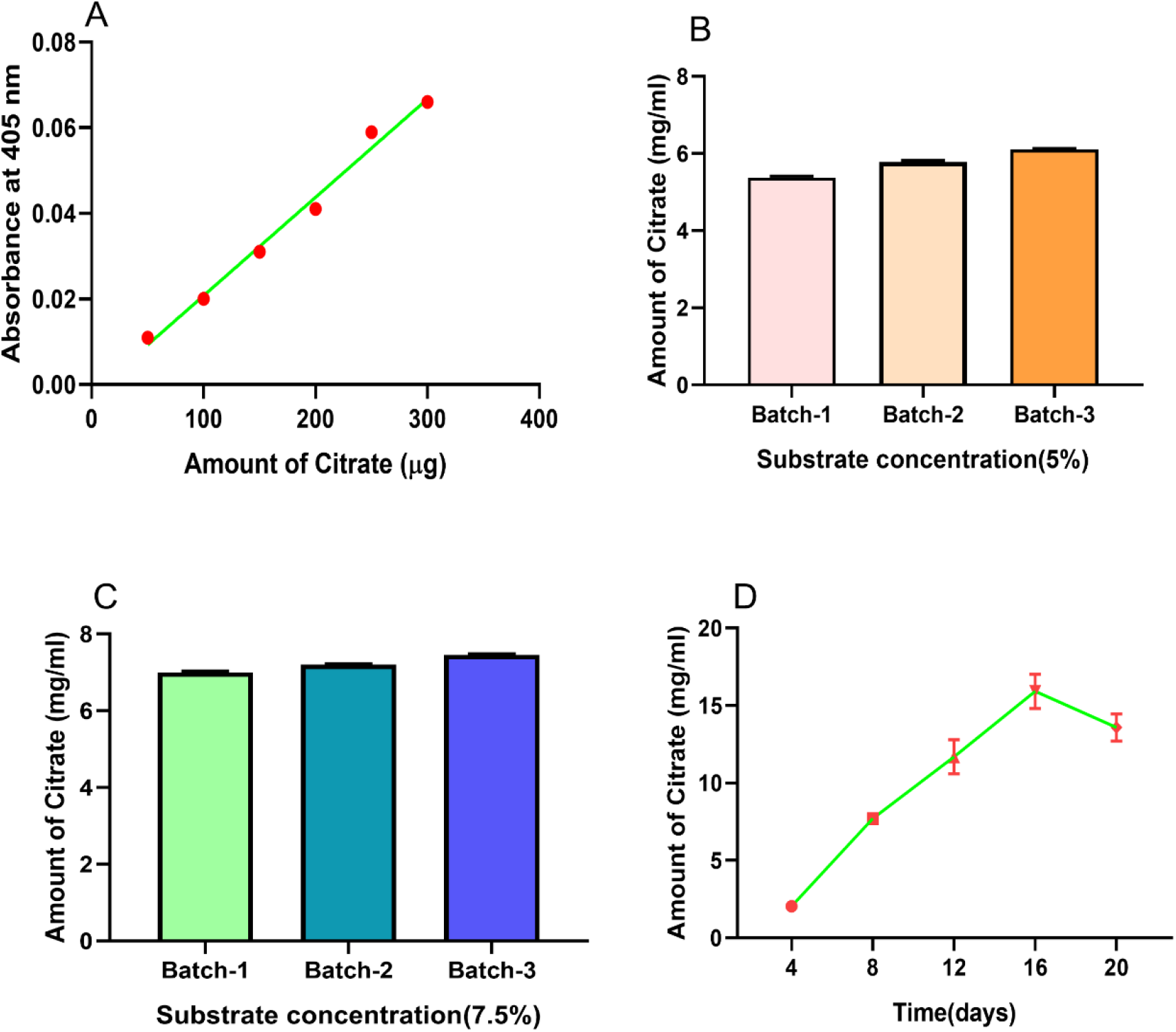
Standardization of Citric acid production using molasses as a raw material. **(A) shows the** titration data of citrate, which were generated by Marier-Boulet method using a standard citrate solution. (B) shows the amount of citrate production in different batches of submerged fermenter using either 5% or (C) 7.5% of molasses as a source of initial substrate. The amount of citrate was determined by non-linear regression method. And (D) shows the optimum duration to produce the maximum amount of citrate by the submerged fermentation using cane molasses as a raw material. Data shows as mean ± standard deviation. N=3.

### Maximum production limit of citrate in submerged fermentation

To observe the effect of pretreatment with potassium ferrocyanide, 5% molasses media was inoculated with inoculum size 2%, initial pH 5.6 for 10 days fermentation. The result had shown that maximum yield of citric acid was observed in a pretreated batch (12.49 mg/ml) compared with untreated batch of fermentation (Figure 4A). *Aspergillus niger* F21 isolate (4% inoculum size) was used for citric acid production using 10% sugarcane as a substrate with the above-mentioned optimum pH and temperature. Maximum citric acid was found 28.25 mg/mL after 16 days fermentation (Figure 4B).

**Figure 4:**
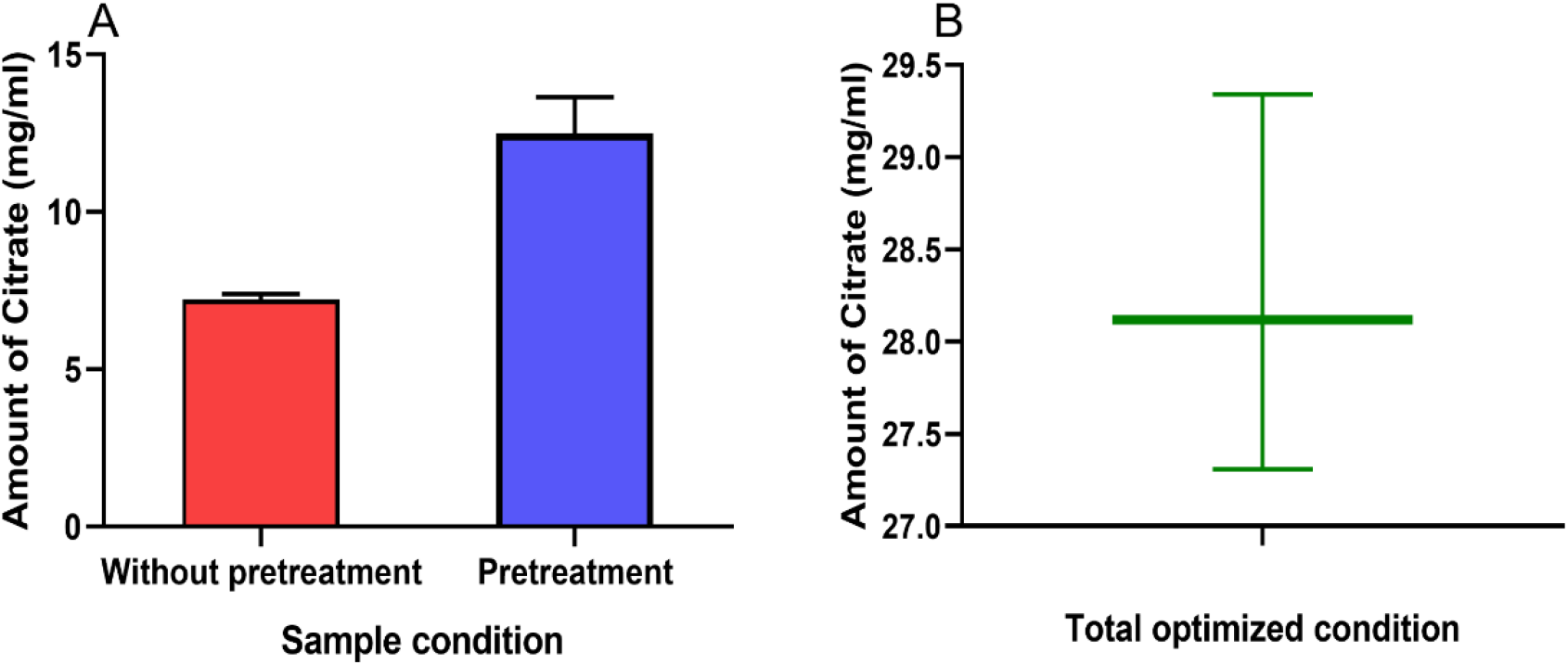
Citrate production limits in cane molasses and *A. niger* containing submerged fermentation. (A) shows the amount of citrate production with/without pretreated cane molasses. (B) shows the highest amount of citrate production using optimized amount of cane molasses, temperature, pH, and inoculum of *A. niger*. Data shows as mean ± standard deviation. N=3.

## Discussion

Citric acid is one of the most demanding organic ingredients because it is use in 70% food-additives, 12% pharmaceuticals and 18% other industrial products (Dhillon et al., 2011). It can be derived from lemon, lime and oranges as a natural source (Penniston et al., 2008). In addition, citrate can be produced synthetically by chemical reaction, but microbial fermentation met 99% of citric acid requirements globally (Kuforiji et al., 2010). Hence this study has adopted the submerged fermentation technique, which is less sensitive to medium composition, allowing a broad range of substrates with flexible substrate control. In addition, this technique is cost-effective with minimum risk of batch contamination and high yielding capacity (Max et al. 2010). Besides, this also allow to use different microbes, including bacteria and fungi to produce citric acid. Current study has utilized *Aspergillus niger*, which is superior to other microbes particularly for citric acid production because its high yielding capacity using cheap raw materials like cane molasses as a source of carbohydrate (Alhadithy, 2020; Show et al., 2015).

Apart from carbohydrate, cultural conditions of *Aspergillus niger* are crucial factors for submerged fermentation-based citrate production (Çevrimli et al., 2009). In this study, the submerged fermentation has adopted to produce citrate because its cost-effective implementation and reasonable reproducibility. However, other parameters for this process have optimized to produce to obtain maximum amount of citric acid. In addition, we also demonstrated that yielding of citrate increased by a decrease of the pH value, and the sugar content in the solution. In this study fermentation conditions and *A. niger* F21 isolates with pH 6.0 appears to be the best initial pH for maximum production of citrate (28.25 mg/ml). Overall, these findings were corroborated with previous findings, where an initial pH 5.5 contributes to the maximum level of citric acid biosynthesis (Ikram-ul et al., 2004; Sikander et al., 2002). However, Khurshid et al., demonstrated that a higher pH causes accumulation of oxalic acid and a low, or high pH causes negative impact on citric acid production due to the release of toxic ions (Khurshid et al., 2024). Furthermore, Dashen and colleagues demonstrated that pH of the medium changes constantly due to microbial metabolic activities, primarily due to the secretion of organic acids such as citric acid, as well as the unwanted gluconic and oxalic acid (Dashen et al., 2014). We also observed that the pH of the fermenter is critical during sporulation of *A. niger* and citrate production.

During the germination stage, the germinating spores absorb ammonia and release protons, increasing the acidity of the medium and favoring citric acid production.

In addition, it is reported that 25-30°C is optimum for citrate production by submerged fermentation. To enhance citrate production wild type strain of *A. niger* needs to acclimatize as an industrial strain, which is yet to establish in our study. This is mostly done through strain improvement process. Moreover, mutagenesis can be applied to improve citric acid producing *A. niger* through inducing the parental strains. For example, Mutagens including gamma radiation, ultraviolet radiation and chemical mutagens are applied to induce mutation. Besides, ultraviolet and chemical mutagens-based combined method produces citrate hyperproducer strain, reviewed in (Chandra et al., 2021). The passage and single spore techniques are used for selection. The passage method is preferred because organic acids (oxalic and gluconic acids) and mineral acids simulate the presence of citric acid in the single-spore method (Soccol et al., 2006). Moreover, this study also optimized the spectrophotometry method to measure the level of citrate in each batch of fermenter, which is reproducible, less time consuming and cost-effective.

## Acknowledgements

The work is supported by two research grants funded by the Ministry of Education, Bangladesh and the Ministry of Science and Technology, Bangladesh. Fungal culture has supported by Molecular Biology lab, department of Biochemistry and Molecular Biology, University of Dhaka, Bangladesh.

## Author contribution

Alamin and RA have equally conducted and analyzed experiments. TNI reviewed the manuscript. MM and MI designed experiments and reviewed analysis with Alamin and RA. MM wrote the manuscript with input of all authors.

## Notes

### Competing Interest Statement

The authors have declared no competing interest.

